# A multi-organ spatial metabolomic atlas of exercising mice reveals neuronal Complex I as a convergent and sufficient axis for tau pathology reduction in PS19

**DOI:** 10.64898/2026.06.16.732673

**Authors:** Terrymar Medina, Sadi Quiñones, Zizhen Liu, Lei Wu, Borhane Ziani, Cameron Shedlock, Xin Ma, Alison Ryan, Reece Larson, Charles M. Soto, Vicenzo Barco-Caiaffa, Ann Titus, Kangjin Wong, Nidhi Rao, Sophia Florea, Miguel A. Gutierrez-Monreal, Gordon S. Mitchell, Craig W. Vander Kooi, Matthew S. Gentry, Navdeep S. Chandel, Karyn A. Esser, Ramon C. Sun

**Author notes:** These authors contributed equally: T.M., S.Q., Z.L. Senior author and Lead contact: R.C.S.

## Abstract

We constructed a spatially resolved metabolomic atlas of long-term exercise across six major organs in wild-type mice: brain, heart, lung, liver, kidney, and skeletal muscle, cataloguing 224 metabolic features and revealing coordinated inter-organ remodeling. Surprisingly, the brain showed particularly pronounced region-specific adaptation. Because pathological tau associates with synaptic mitochondria from early stages of tauopathy, we extended this multi-organ spatial metabolomic approach to PS19 mice and found that exercise reduced over 70% of observable tau pathology in PS19 hippocampus and restored the mitochondrial-related metabolome. Integrated proteomic and spatial metabolomic analyses identified NADH dehydrogenase Complex I as the convergent node. To test this finding biologically, we expressed the yeast NADH dehydrogenase, Ndi1, in PS19 neurons in the absence of exercise. This increased cerebral antioxidants, restored shuttle-linked metabolites, and reduced tau pathology. Increasing NADH dehydrogenase activity through NDI1 reproduces the core anti-tau and metabolic effects of exercise. These findings provide a molecular mechanism for how exercise may prevent or slow tau pathology accumulation, complementing the human-cohort literature linking exercise to delayed cognitive decline.

## Introduction

Regular physical exercise induces coordinated physiological adaptations across nearly every major organ system and is among the top non-pharmacological interventions to combat non-communicable disease and promote healthy aging (1–7). Work over the past decade has established that exercise can elicit systemic effects via circulating molecules termed ‘exerkines’ (8), as well as exercise-inducible metabolites with systemic effects on feeding and energy balance (9, 10). These ‘exerkines’ are secreted from muscle, liver, bone, and adipose, and are responsible for many of the health benefits observed with exercise.

Among the organs influenced by exercise, the brain is particularly responsive. Regular physical exercise protects cognitive function across the lifespan (1, 11–13). Voluntary wheel running in rodents remodels the adult hippocampus, promoting neurogenesis, synaptic plasticity, and spatial memory (11, 14). Endurance training in older adults increases hippocampal volume and protects against age-related cognitive decline (12, 15). In humans at risk of Alzheimer’s disease (AD), regular physical activity is associated with slower tau accumulation, lower neurofilament light chain, and preserved cognition (16–18). Lifestyle trials including the U.S. POINTER study now provide randomized evidence that structured exercise can meaningfully alter dementia risk (19, 20). Single-cell and spatial omics studies in mouse models of brain aging and Alzheimer’s disease have begun resolving the cellular and regional architecture of exercise responses in the diseased brain (21, 22). Tau pathology is the principal neuropathological and imaging correlate of synaptic loss and cognitive decline across AD and primary tauopathies (23). Tau is predominantly a neuronal protein, and pathological tau aggregates initiate within neurons and propagate trans-synaptically along connected circuits in AD (24, 25), locating the disease at the level of neuronal cell biology. Yet despite consistent evidence that exercise prevents or slows cognitive decline in at-risk populations, the molecular mechanisms through which exercise engages the diseased brain, particularly in primary tauopathy, remain poorly defined (1, 26). Defining these mechanisms in animal models that permit early intervention is essential for developing pharmacologic strategies for patients unable to exercise.

In mouse tauopathy models, pathological tau associates with synaptic mitochondria as early as 3 months of age, coinciding with synaptic energetic dysfunction and excitatory synapse loss (27). Mitochondrial respiration, biogenesis, and dynamics are similarly impaired well before overt neurodegeneration (28, 29). Brain bioenergetic dysfunction is another modifier of dementia. Hippocampal glucose hypometabolism precedes clinical cognitive decline by several years on FDG-PET (30), and inherited and age-dependent bioenergetic decline is positioned upstream of both Aβ and tau pathology in the mitochondrial cascade hypothesis (31, 32). This places brain energy metabolism among the most actionable axes of dementia therapy (33). Among bioenergetic defects in AD, the most consistent is electron transport chain impairment. Decrements in Complex I and cytochrome c oxidase activity have been documented in AD platelets and post-mortem cortex since the early 1990s (34, 35) and are featured prominently in neuropathological surveys of mitochondrial dysfunction (36–38); more recently, in vivo PET imaging has linked Complex I dysfunction to neurodegeneration and cognitive decline in AD (39). Neurons are especially sensitive: synaptic transmission consumes a disproportionate share of neuronal ATP, and sustained oxidative phosphorylation requires constant re-oxidation of cytosolic NADH to fuel the electron transport chain via Complex I (40, 41). Complex I activity is therefore crucial for normal brain function. The malate–aspartate shuttle is a major route for this cytosol-to-mitochondria redox transfer, while Complex I provides the principal mammalian NADH:ubiquinone entry point into the ETC.

The yeast alternative NADH dehydrogenase Ndi1 provides a genetic strategy to bypass mammalian Complex I-mediated NADH:ubiquinone electron entry. Ndi1 oxidizes mitochondrial NADH and transfers electrons to ubiquinone without proton pumping (42, 43). Ndi1 has rescued Complex I deficiency in Drosophila neurons (44), dopaminergic neurons in MPTP-induced parkinsonism (45), retinal ganglion cells in glaucoma (46), and amyloid-induced AD pathology in mice (47). Ndi1 thus provides a unique tool to test the phenotypic consequences of restoring electron flow in biological systems.

Here, we leveraged matrix-assisted laser desorption/ionization mass spectrometry imaging (MALDI-MSI) (48, 49) to generate a multi-organ spatial metabolomic atlas of the long-term exercise response in wild-type mice after 3 months of voluntary wheel running. To study the neurobiological effects of exercise, we examined the brain in much greater detail. In wild-type animals, exercise reshaped the brain metabolome across multiple regions. In PS19 mice, exercise remodeled the disease metabolome and reduced tau pathology vs sedentary littermates. Integrated metabolomics and whole brain proteomics converged on Complex-I–coupled electron transport and the malate-aspartate shuttle as the convergent metabolic signature associated with the exercise response. To test whether restoration of neuronal Complex I electron transport is sufficient to recapitulate the beneficial effects of exercise on the brain, we delivered AAV-SYN1-NDI1 via intracerebral ventricular injection to sedentary PS19 mice. NDI1 treatment increased antioxidants, restored the malate-aspartate shuttle metabolites, and reduced tau pathology – similar to what was observed by exercise. Together, these results highlight the therapeutic potential of exercise mimetics and position neuronal Complex I as a candidate bioenergetic node engaged by the exercise response.

## Results

### Multi-organ spatial metabolomic atlas of long-term exercise

To investigate the multi-organ metabolic effects of long-term exercise in mice, we used the voluntary wheel running paradigm, in which C57BL/6J (WT) mice were given continual, free access to a running wheel from 3 months-, to 6 months of age. Control mice were placed in similar housing conditions but did not receive a running wheel. Daily running activity was recorded to verify consistent exercise behavior (Fig. S1, table S1). Following 3 months of exercise training, the running wheels were locked for 3 days to eliminate acute exercise effects on the metabolome, after which mice were euthanized and their organs harvested for untargeted spatial metabolomic analysis via MALDI-MSI (Fig. 1, A and B). In total, we collected 6 organs per mouse: brain, heart, lung, liver, kidney, and skeletal muscle (quadricep). Each of these whole organs was thinly sectioned and scanned at 50 µm resolution to enable organ-, and subregion-, specific metabolomic analysis. After quality control (QC) filtering and batch processing of the data (Materials and methods, Fig. S2), we identified a total of 224 unique metabolic features with varying organ-specific detection (Fig. S3). Next, we performed unsupervised, integrated spatial clustering to identify distinct metabolic zones within each organ (Fig. S4, A, G, L, Q, V; Materials and methods). When assessed against H&E staining of the sequential tissue section, these organ-specific metabolic zones appear to correspond to distinct tissue architecture (Fig. S4, C, I, N, S, X). To that point, within the heart, we identified two main clusters that strongly map to the cardiac tissue and cardiac cavity (the blood-filled chambers of the heart), respectively (Fig. S4, A and B). Given the physiological difference between the cardiac tissue and cardiac cavity, these were analyzed as unique organs in downstream analysis.

**Fig. 1.**
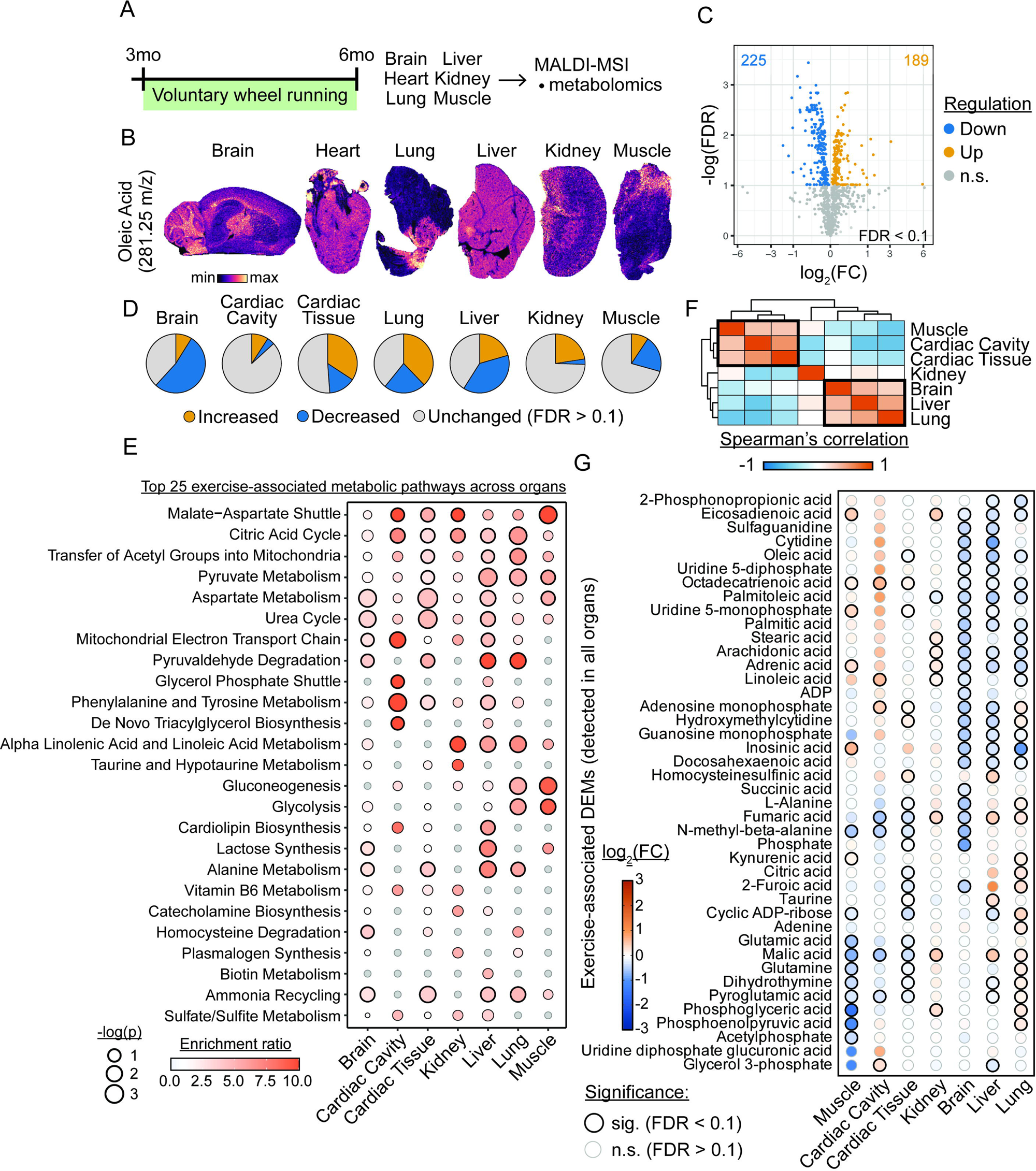
Multi-organ spatial metabolomic atlas of chronic exercise. (A) Experimental design schematic. n = 6 organs per animal, N = 3 animals per group. (B) Representative ion images (Oleic acid, 281.25 m/z) from each organ. (C) Volcano plot of all differentially expressed small molecule features across each organ. (D) Pie charts showing relative fraction of small molecules with significant change vs no change, per organ. (E) Dotplot of top 25 exercise-associated metabolic pathways across organs, ranked by enrichment ratio. Size = significance, fill = enrichment ratio. Grey fill = not enriched/NA. (F) Spearman correlation matrix demonstrating inter-organ dynamics. Correlation based on log2FC (exercise vs control). (G) Dotplot of exercise-associated changes to small molecules shared across organs. Bold dots = significant changes. Significance throughout (A–G) determined by Wilcoxon rank sum with Benjamini–Hochberg FDR correction (FDR < 0.1 unless noted otherwise); N = 3 mice per group.

To identify the effect of long-term exercise on these organs, we ran whole organ and cluster-specific differential expression (DE) analysis (Fig. 1C, fig. S4, D, E, J, O, T, Y, and Materials and methods). In support of our spatial approach, cluster-specific DE analysis revealed subregion-specific metabolic changes that were not detected in whole organ DE analysis (Fig. S4, F, K, P, U, Z). Alanine, shown to protect against chronic kidney disease (50), was found to be increased in a metabolic subregion of the kidney (Fig. S4). Meanwhile in the cardiac cavity, we detected an increase in n-acetyl-l-phenylalanine – an n-acetylated exerkine similar to Lac-Phe (51) (Fig. S4, AA and AB).

Since our goal was to understand multi-organ regulation, and we found very few (<1%) features with opposing responses among organ clusters, we restricted our downstream analysis to whole organ DE. This revealed 414 total organ-feature changes (225 decreased and 189 increased) across the 6 organs, drawn from the 224 unique metabolites detected (Fig. 1C; Materials and methods).

After normalizing each organ’s total DE features to its total detected library, we found that the highest ‘exercise training-responders’ were the brain, lung and liver (Fig. 1D). Correlation analysis on the exercise-associated DE features of each organ (Materials and methods) revealed that the brain, lung, and liver were significantly positively correlated, while the muscle, cardiac cavity, and cardiac tissue were inversely correlated – suggesting inter-organ metabolic remodeling consistent with the tissue-specific exercise responses catalogued by MoTrPAC (7, 52) (Fig. 1, F and G). The kidney did not exhibit positive correlation with any other organ (Fig. 1, F and G).

To contextualize overall metabolic changes to each organ, we performed metabolic pathway analysis (Materials and methods) on all DE features, including organ-specific features (Fig. 1E). This analysis largely revealed metabolic pathways known to be regulated by exercise (e.g., ‘Gluconeogenesis,’ ‘Citric Acid Cycle,’ ‘Mitochondrial Electron Transport Chain’). Plotting the top 25 most enriched pathways across organs exhibited substantial organ-dependent changes (Fig. 1E). For example, the liver exhibited significant enrichment of the ‘Cardiolipin Biosynthesis’ pathway, a critical effector of thermogenesis linked to insulin sensitivity in humans (53). Other organ-specific enriched metabolic pathways include: ‘Homocysteine Degradation’ in the brain, associated with dementia and stroke (54, 55), and ‘Glycolysis’ and ‘Gluconeogenesis’ in the lung and muscle (Fig. 1E).

Taken together, these results demonstrate a spatially resolved atlas of exercise-associated metabolic remodeling and reveal both organ-specific responses and coordinated metabolic signatures across organs.

### Brain region-specific metabolic changes associated with long-term exercise

To determine whether exercise training effects are heterogeneous across brain regions, we started by annotating 6 distinct regions including cortex (CTX), hippocampus (HP), white matter (WM), thalamus (TH), striatum (ST), and cerebellum (CB) (Fig. 2A; Materials and methods).

**Fig. 2.**
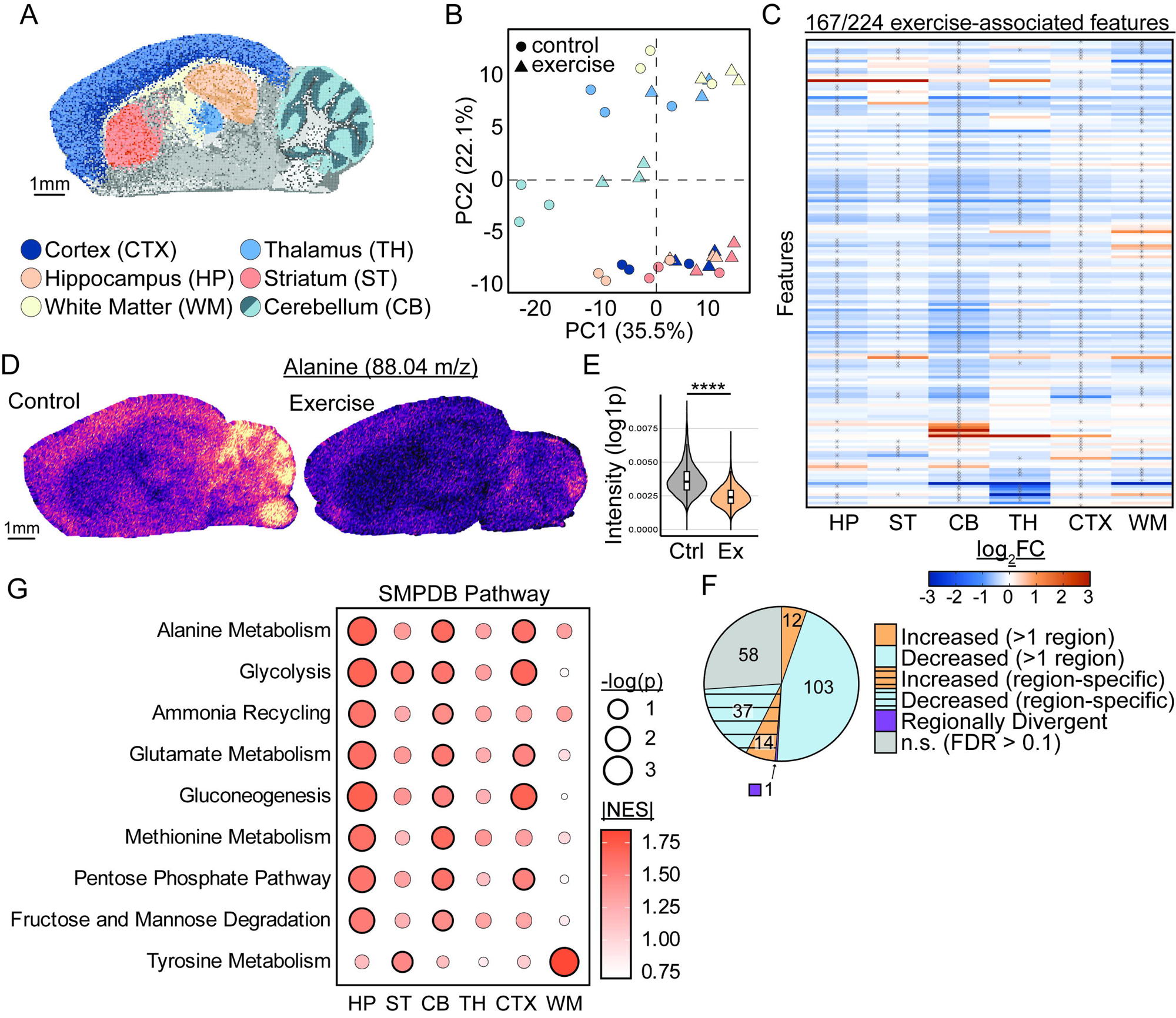
Brain region-specific metabolic changes associated with chronic exercise. (A) Representative spatial mapping of brain region annotations used for downstream region-based analysis. (B) Principal component analysis (PCA) scatter plot showing each annotated brain region across treatments (control, exercise). Each dot represents the mean aggregate for that region in a single animal. Color = region. Shape = treatment (circle = control, triangle = exercise). (C) Heatmap of 167 small molecule features with significant exercise-associated change, in at least 1 region (FDR < 0.1). (D) Representative ion images (Alanine, 88.04 m/z) from control and exercise. (E) Violin plot quantification for alanine in (D). Violin represents per-pixel distribution for each representative section. **** p < 0.0001 by Wilcoxon rank sum with Benjamini–Hochberg FDR correction. Sample size = 19,125 unique pixels (Control); 18,730 unique pixels (Exercise). (F) Pie chart breakdown of region-specific vs non-region-specific exercise-associated changes. (G) Dotplot of top 9 exercise-associated metabolic pathways, ranked by normalized enrichment score (NES) and significance. Size = significance, fill = absolute value of NES. Scale bar in (A) and (D) = 1 mm.

Next, we ran principal component analysis (PCA) on all 224 detected features to gain a global perspective of the exercise effect across regions studied here (Fig. 2B). Consistent with brain region-specific metabolic studies, the brain regions (regardless of treatment) exhibited substantial region-driven spread across PC2 (22.1% variance). However, the exercise-driven PC1 (35.5% variance) exhibited no region-specific differences (Fig. 2B), suggesting a generalizable exercise effect on brain metabolism, at least from a high-level perspective.

To deeply probe these regional differences, we performed region-specific DE analysis and quantified the number of features conserved (>1 region), divergent, region-specific, or non-significant (FDR > 0.1) (Fig. 2, C and F). Many features were conserved across brain regions, with only 1 feature exhibiting significant divergent response: a hexose-6-phosphate (m/z 259.02, consistent with fructose-6-phosphate but other isomers cannot be excluded) (Fig. 2F). Of the 51 region-specific DE features observed, many were qualitatively conserved in other regions. Alanine, for example, displayed pronounced region-specific changes between control and exercise (Fig. 2, D and E). Overall, this analysis supports the idea that many metabolic effects of exercise are generalizable across brain regions. However, whereas the features themselves do not display much regional heterogeneity, significant differences in the magnitudes of feature changes were observed (Fig. 2C).

To quantify pathway-level differences between brain regions, we performed metabolite set enrichment analysis (MSEA; Materials and Methods) using ranked log fold changes, to enable comparison of pathway enrichment strength across regions with shared metabolite features. In this analysis, many top exercise-associated metabolic pathways were significantly perturbed across 3 major regions – HP, CB, and CTX (Fig. 2G). Conversely, WM exhibited distinct and preferential enrichment of only 1 of these top pathways – ‘Tyrosine Metabolism.’ Taken together with the feature-level coherency observed, these pathway results may suggest that, while large divergent responses are minimal, exercise produces variable magnitudes of change in the most notable regions: hippocampus, cortex, and cerebellum.

### Exercise reduces hippocampal p-tau and remodels the PS19 brain metabolome

Phosphorylated tau (p-tau) is a defining pathological feature of several neurodegenerative diseases, most notably Alzheimer’s disease (AD). Emerging studies on rodent models and humans with preclinical AD demonstrate a significant relationship between long-term exercise and reduced tau burden (16–18, 56–59) – a characteristic that correlates with cognitive decline. However, the molecular drivers of these effects are unknown. To address this gap, we utilized the P301S (PS19) tauopathy mouse model, which develops hippocampal p-tau pathology, to determine whether 3 months of voluntary wheel running could recapitulate tau reduction in the brain, and to identify potential therapeutic nodes for tau reduction based on downstream exercise effects. WT and PS19 mice were placed in long-term wheel running housing or control (no wheel) conditions, as described above, from ages 3 to 6 months. Total running and body weight were unchanged across genotypes (Fig. S1). After exercise (or non-exercise), whole brains were harvested for: immunofluorescence (IF) staining (anti-p-tau), MALDI metabolomics, and untargeted proteomics (Fig. 3A).

**Fig. 3.**
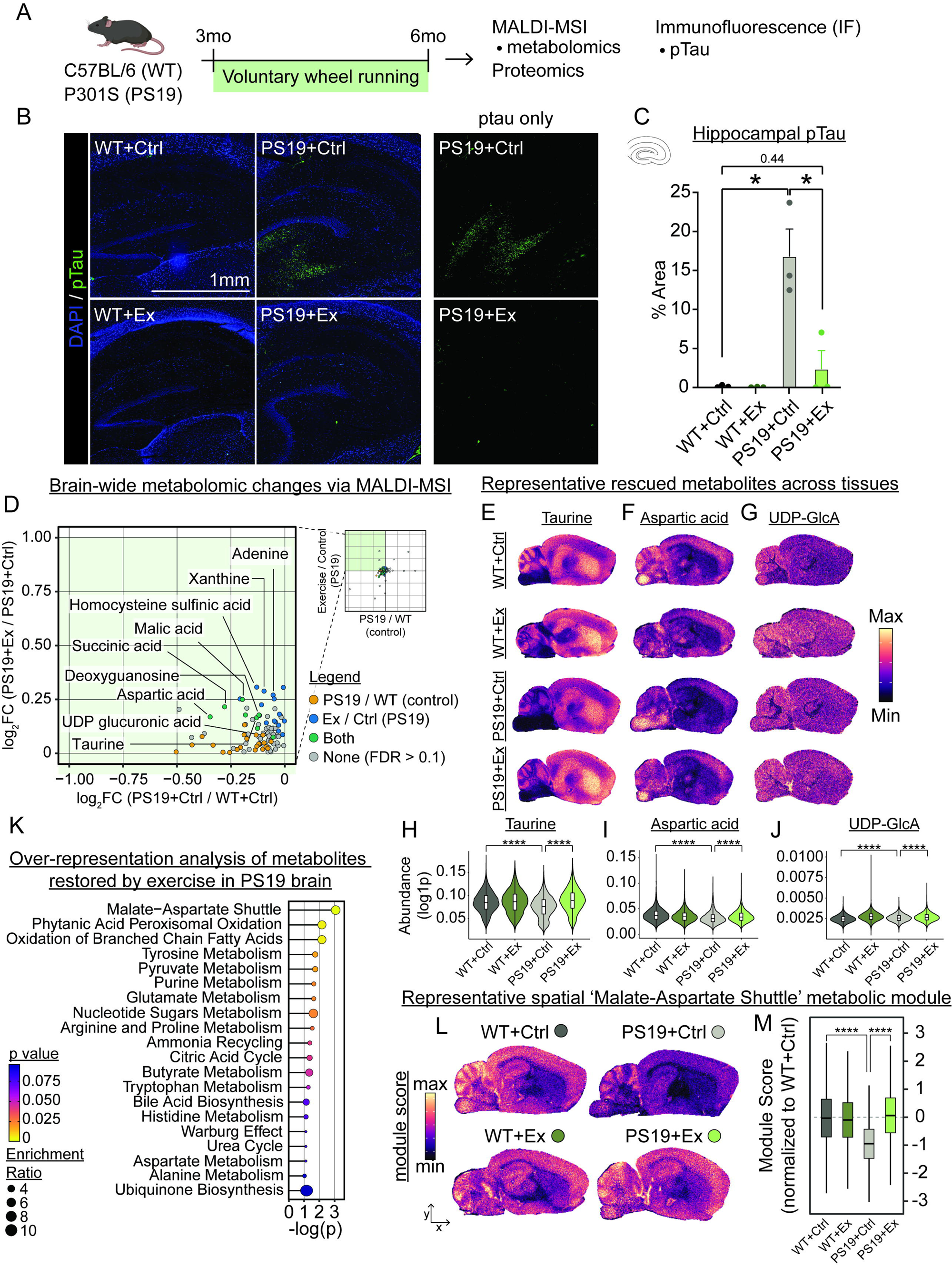
Chronic exercise induces metabolic remodeling associated with attenuation of phosphorylated tau (p-tau) in PS19 mice. (A) Experimental design schematic. n = 6 organs per animal, N = 3 animals per group. Created with Biorender. (B) Representative immunofluorescent (IF) images of hippocampal p-tau across groups. p-Tau only images of PS19 groups are shown on (right). (C) Quantification of relative p-tau abundance across groups. Error bars = SEM. * p < 0.05 by Welch’s t-test. Sample size = 3 mice per group. (D) Scatterplot showing the brain-wide PS19-depleted, exercise-enriched small molecule features. Colored by significance within each comparison. (E to G) Representative brain ion images from control and exercise. Taurine (m/z) (E), Aspartic acid (m/z) (F), and UDP-GlcA (m/z) (G). (H to J) Violin plot quantification for each representative ion image in (E to G): taurine (H), aspartic acid (I), and UDP-GlcA (J). Violins represent per-pixel distribution for each representative section. **** p < 0.0001 by Wilcoxon rank sum with Benjamini–Hochberg FDR correction. (K) Dotplot showing over-representation analysis results driven by rescued small molecule features identified in (D). Size = enrichment ratio, fill = significance. (L) Representative spatial mapping of ‘Malate-Aspartate Shuttle’ module scores, across groups. (M) Per pixel quantification of module scores based on representative brains in (L). **** p < 0.0001 by Wilcoxon rank sum with Benjamini–Hochberg FDR correction. Sample size for per pixel quantification in (H to J and L): 19,125 unique pixels (WT+Ctrl); 18,730 unique pixels (WT+Ex); 20,184 unique pixels (PS19+Ctrl); 19,795 unique pixels (PS19+Ex). Scale bars in (B, E–G, L) = 1 mm.

IF against phospho-Tau Thr231 (AT180 epitope) was used to determine if exercise reduced p-tau in PS19 mouse brains. Quantitative analysis was focused on the hippocampus given its strong enrichment of p-tau at this age (Fig. SX) and the central role of the hippocampus in AD. As expected, substantial enhancement of p-tau immunoreactivity was observed in PS19 hippocampus vs WT (Fig. 3 B and C). Further, exercise decreased p-tau levels in PS19 mice (85% reduction, P < 0.05, PS19+Exercise vs PS19+Control) (Fig. 3 B and C). Next, brain-wide MALDI metabolomic data were analyzed across genotypes (WT, PS19) and treatment (Exercise, Control) to test whether exercise can attenuate metabolic dysfunction in PS19. Since prior MALDI analysis revealed largely coherent changes to brain metabolism in exercise-induced responses, subsequent MALDI DE analyses were run using an integrated brain-wide approach (Materials and methods).

To interrogate exercise-associated restoration of PS19-deficient metabolism, we visualized significant PS19 genotype-associated metabolic changes (PS19+Control vs WT+Control, FDR < 0.1) vs exercise-associated metabolic changes to PS19 mice (PS19+Exercise vs PS19+Control, FDR < 0.1) (Fig. 3D). We defined a feature as ‘rescued’ when it was significantly altered by PS19 genotype (PS19+Control vs WT+Control) and significantly reversed in direction by exercise (PS19+Exercise vs PS19+Control). From this cross-analysis, 8 metabolite features met these rescue criteria, decreased by genotype and restored by exercise (Fig. 3D). These metabolite features included amino acids (e.g., taurine and aspartic acid) and nucleotide sugars (e.g., UDP-GlcA) (Fig. 3 E – G). Individual representative spatial heatmaps and pixel-level violin plots of each feature highlight the inherent variability in expression across regions (Fig. 3 E – J).

Next, we ran metabolic over-representation analysis (ORA) of the 8 features, combined with 16 metabolite features that were significantly increased by exercise in PS19, but only insignificantly (but qualitatively) decreased by genotype, to determine whether these ‘rescue’ features corresponded strongly to any given metabolic pathway (Materials and methods). ORA revealed 3 significant pathways (P < 0.01), each of which appears to converge on mitochondrial metabolism – ‘Malate-Aspartate Shuttle’, ‘Phytanic Acid Peroxisomal Oxidation’, and ‘Oxidation of Branched Chain Fatty Acids’ (Fig. 3K). The most significantly over-represented pathway, ‘Malate-Aspartate Shuttle’ (P < 0.001), was visualized and quantified in representative brain sections as a metabolic module – highlighting the broad effects on this pathway (Fig. 3 L and M; Materials and methods).

### Metabolomic and proteomic analysis of rescue features converge on mitochondrial electron transport chain (ETC)

Next, we analyzed whole brain untargeted proteomic data from each genotype and treatment group. As for metabolomics, we visualized significant proteomic DE analysis results from the genotype effect (PS19+Control vs WT+Control, FDR < 0.05) vs exercise-associated protein changes in PS19 mice (PS19+Exercise vs PS19+Control, FDR < 0.05) (Fig. 4A; Materials and methods). Hundreds of proteins were identified that were significantly rescued by exercise in PS19, including proteins aberrantly increased and decreased in PS19 (Fig. 4A). We quantified the number of ‘rescue’ proteins against the total number of proteins affected by PS19 genotype and found that ∼28% of all genotype-associated proteins were reversed by exercise (Fig. 4B).

**Fig. 4.**
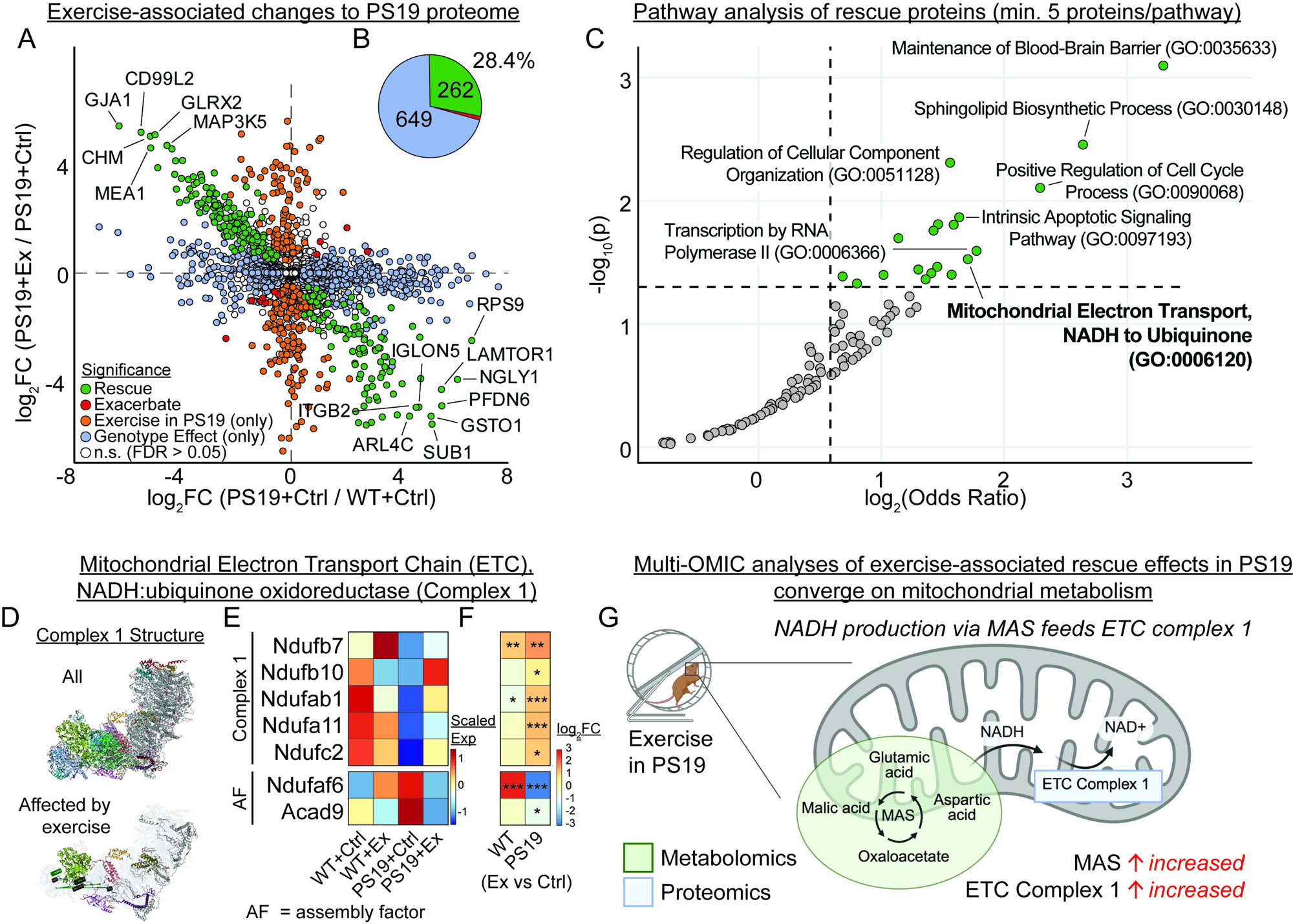
Metabolomic and proteomic analysis of rescue features converge on mitochondrial electron transport chain (ETC). (A) Scatterplot of differentially expressed proteins across genotype (PS19+Ctrl vs WT+Ctrl) (x-axis) and treatment (PS19+Ex vs PS19+Ctrl) (y-axis). Colored by significance within each comparison. Sample size: N = 3 mice per group. (B) Pie chart showing percent of all PS19-associated proteins with significant exercise-associated rescue. (C) Scatterplot of enriched pathways based on ‘rescued’ proteins in (A). Minimum of 5 proteins per pathway. Colored by significance + odds ratio threshold. (D) Cryo-EM structure of Complex I (PDB: 7QSK; top), colored by exercise effect (bottom). (E) Heatmap of average Complex I subunit and assembly factor (AF) protein expression values per group. (F) Heatmap of log2FC (Ex vs Ctrl) per genotype for proteins in (E). *** p < 0.001, ** p < 0.01, * p <0.05 by empirical Bayes moderated DE analysis with Benjamini–Hochberg FDR correction. (G) Schematic illustration of the convergent mitochondrial mechanism observed between metabolomic and proteomic analysis of exercise in PS19. Metabolomics (green) depict increase in malate-aspartate shuttle observed in Fig. 3K. Proteomics (blue) depict increase in ETC Complex I observed in (C to F). Created using Biorender.

To contextualize proteomic rescue, we ran pathway analysis on the 262 rescue proteins identified in (A) (Materials and methods). From those rescued pathways, we identified 19 significant pathways, including ‘Maintenance of Blood-Brain Barrier’, ‘Intrinsic Apoptotic Signaling Pathway’, and ‘Mitochondrial Electron Transport, NADH to Ubiquinone’ (Fig. 4C). These represent rescued pathways that may serve as potential candidate mechanisms underlying exercise benefit to tauopathy.

In parallel to our DE analyses, we applied weighted gene co-expression network analysis (WGCNA) to our full proteomic dataset to identify naturalistic, coordinated protein networks associated with exercise rescue in PS19 (Fig. S5A). WGCNA revealed multiple modules whose expression patterns tracked genotype and treatment status, including a brown module that was reduced in PS19 and restored by exercise, and a red module that exhibited an opposite pattern (Fig. S5 B to D). The brown module includes proteins involved in ubiquitin-dependent proteostasis, such as Psmc3, Psmc5, Psme1, Uchl5, and Rad23b (Fig. S5E). Notably, the module was further anchored by several highly connected hub proteins (as defined by kME > 0.8), including Glrx2, Stk11/LKB1, Lamtor3, C9orf72, and Cmpk2 (Fig. S5F). These hub proteins are functionally linked to mitochondrial redox homeostasis (60, 61), cellular energy sensing via AMPK (62), lysosome-associated nutrient sensing (63), and innate immune-metabolic regulation (64). Together, this analysis implicates coordinated regulation of mitochondrial homeostasis, redox balance, proteostasis, and cellular energy sensing as central features of the exercise-restored network – complementary to our DE pathway analysis that independently identified restoration of mitochondrial electron transport.

Mitochondrial electron transport chain (ETC), NADH:ubiquinone oxidoreductase, also known as ETC Complex I, is comprised and regulated by core structural components and dedicated assembly factors (Fig. 4D, fig. S6A). To determine the directionality of exercise effects on Complex I, we plotted the group averages and the log2FC (Exercise vs Control, per genotype) of each significantly regulated (FDR < 0.05, PS19+Exercise vs PS19+Control) Complex I structural component and assembly factor (Fig. 4 E and F). Broad exercise-associated reversal of the PS19-associated Complex I proteomic signature was detected, defined by an exercise-associated increase in PS19-deficient core structural components, and reversal of PS19-associated increases in assembly factor expression, but no apparent change to proton pump machinery (Fig. 4 E and F, fig. S6A). The independent metabolomic and proteomic analyses converge on the entry point of the ETC: NADH oxidation by Complex I downstream of MAS-supplied NADH (Fig. 4G). This positions Complex I restoration as a candidate mechanism underlying the exercise response in PS19 brain.

Finally, we leveraged our rescue metabolites (n=24) and rescue proteins (n=262) to perform integrated multiOMIC pathway analysis (Materials and methods, fig. S7A). We tested against the KEGG (Kyoto Encyclopedia of Genes and Genomes) database, which characterizes pathways based on small molecules (e.g., metabolites) and genes/proteins. Fifteen significant pathways (P < 0.01, enrichment ratio > 2) were revealed in 3 broad categories: neurodegenerative disease (e.g., Amyotrophic lateral sclerosis, Huntington disease, Alzheimer disease), metabolism (e.g., oxidative phosphorylation, thermogenesis, taurine and hypotaurine metabolism), and cell signaling (e.g., tight junction, gap junction, retrograde endocannabinoid signaling) (Fig. S7B). This convergence on canonical neurodegeneration pathways links our rescue signature with established disease mechanisms and supports the multi-OMIC framework.

### Neuron-targeted NDI1 rescues hippocampal p-tau immunoreactivity in sedentary PS19 mice

Complex I is found in neurons and glial cells, yet its assembly, regulation, and expression do not appear uniform across cell types or regions (65, 66). Using rodent brain MERFISH data from Zhang et al. (2023) (67), we observed robust neuron-enriched expression of Complex I genes, including the exercise-associated Complex I components (Ndufb10, Ndufab1, Ndufa11, Ndufb7, Ndufc2, Ndufaf6, Acad9), relative to other cell types (Fig. S6 B to G). This robust neuronal expression persisted at the protein level (Fig. S6D).

To test whether augmenting neuron-specific NADH dehydrogenase activity is sufficient to recapitulate the exercise-associated rescue in PS19, we evaluated 3 orthogonal endpoints: i) hippocampal phosphorylated tau by immunostaining, ii) brain-wide spatial metabolomics by MALDI-MSI, and iii) whole-brain proteomics. Concordant rescue across these readouts would establish that augmenting neuronal Complex I activity is sufficient to reproduce the exercise-associated benefit. To target neuronal NADH dehydrogenase activity directly, we took advantage of the yeast single-subunit NADH dehydrogenase Ndi1, a non-proton-translocating enzyme that can functionally replace Complex I-mediated electron transfer (Fig. 5A). AAVs were delivered to 3-month-old PS19 mice via single intracerebral ventricular (ICV) injection of either neuron-specific NDI1 (AAV9-SYN1-NDI1-eGFP; NDI1) or empty vector (AAV9-CAG-eGFP; control) (Fig. 5B). Co-staining for the pan-neuronal marker NeuN showed that eGFP expression was predominantly detected in NeuN+ neurons, consistent with neuron-enriched Ndi1 expression (Fig. 5C). To mimic the timing of earlier exercise experiments, mice were housed for 3 months without access to running wheels, then brains were harvested at 6 months of age. Our question: can restoration of neuron-specific Complex I activity (electron transfer) replicate the exercise-induced attenuation of p-tau pathology in PS19 brains? We used IF against p-tau in PS19+NDI1 vs PS19+Control brains to quantify relative levels of p-tau immunoreactivity with and without treatment. As before, quantitative analysis was focused on the hippocampus. NDI1-treated brains showed substantial reduction in hippocampal p-tau vs empty vector-treated brains (Fig. 5 D and E), establishing the first of 3 rescue endpoints. Metabolomic and proteomic axes were then assessed in the same animals.

**Fig. 5.**
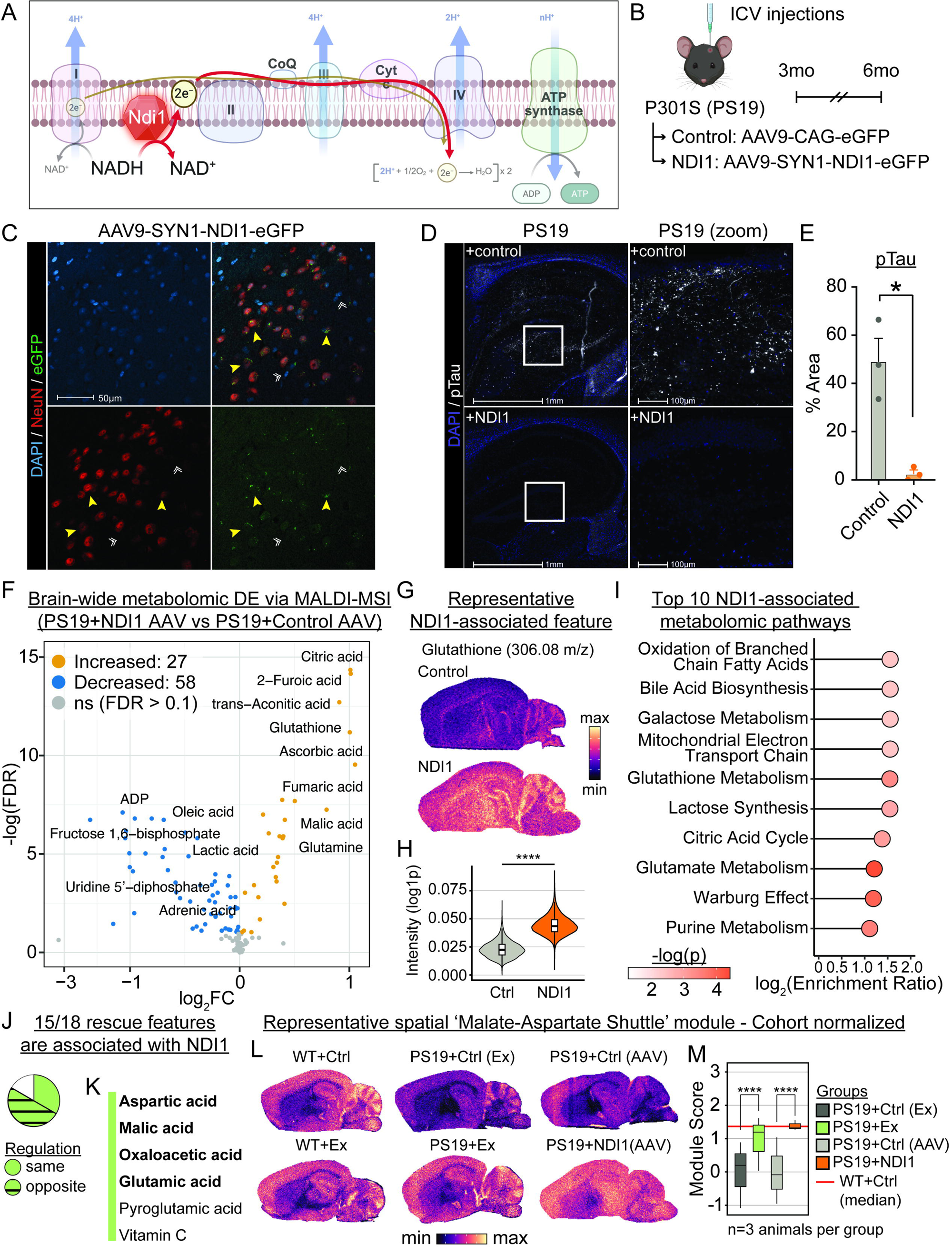
Neuronal NDI1 treatment attenuates p-tau immunoreactivity and induces metabolic remodeling. (A) Cartoon depiction of Ndi1 within the ETC. Red arrows depict Ndi1-mediated NADH dehydrogenase activity and electron transfer. Created using Biorender. (B) AAV experimental design schematic. (C) Representative IF images depicting selective expression of enhanced green fluorescent protein (eGFP) in neurons (NeuN+). Yellow arrows = examples of neurons with eGFP colocalization. White double arrows = examples of non-neurons (NeuN-) with no eGFP colocalization. (D) Representative IF images of hippocampal p-tau. White squares denote location of zoom-in inlets shown on (right). (E) Quantification of relative p-tau abundance across groups. Error bars = SEM. * p < 0.05 by Welch’s t-test. N = 3 mice per group. (F) Volcano plot of brain-wide NDI treatment-associated small molecule features. Colored by significance. (G) Representative ion images (Glutathione 306.08 m/z). (H) Violin plot quantification for glutathione in (G). Violin represents per-pixel distribution for each representative section. **** p < 0.0001 by Wilcoxon rank sum with Benjamini–Hochberg FDR correction. Sample size: 17,178 unique pixels (Control); 18,181 unique pixels (NDI1). (I) Dotplot of top 10 NDI1-associated metabolic pathways, ranked by enrichment ratio. Color = significance. (J) Pie chart showing fraction of exercise-associated features observed in NDI1-associated analysis in (F). (K) List of small molecule features in (I) with same regulation between exercise and NDI1 treatment. Bold features represent malate-aspartate shuttle features. (L) Representative spatial mapping of ‘Malate-Aspartate Shuttle’ module scores, across groups. (M) Quantification of ‘Malate-Aspartate Shuttle’ module scores across all PS19 brains (n=3 animals per group; statistical testing run at pixel-level). Red line = WT+Ctrl median. **** p < 0.0001 by Wilcoxon rank sum with Benjamini-Hochberg FDR correction. Sample size: n > 50,000 unique pixels per group across 3 animals (biological replicates) per group. Scale bars: 50 µm in (C); 1 mm in (D) main panels with 100 µm in (D) zoom-in inlets; 1 mm in (G) and (L).

### NDI1-associated metabolomic and proteomic profiling confirms mitochondrial metabolism remodeling

To address the remaining 2 rescue endpoints, we performed MALDI spatial metabolomics on PS19+Control and PS19+NDI1 brains and whole-brain proteomics from matched samples (Fig. 5 F – L, fig. S8). While the role of Ndi1 in mitochondrial health has been described, less is known of the overall metabolic impact of this treatment in vivo – especially in the context of tauopathy-related neuro-dysfunction.

Using brain-wide metabolomic DE analysis of neuron-directed NDI1 treatment, we identified 85 DE metabolite features associated with NDI1 treatment in PS19 brains (27 increased and 58 decreased, FDR < 0.1) (Fig. 5F). Among the most-affected metabolite features were antioxidants (glutathione, ascorbic acid), TCA cycle intermediates (citric acid, fumaric acid), and glycolysis related molecules (fructose 1,6-bisphosphate, lactic acid) (Fig. 5F), with glutathione shown as a representative example (Fig. 5, G and H). We further characterized these metabolic changes using pathway enrichment analysis and found these NDI1-associated pathways to include, ‘Oxidation of Branched Chain Fatty Acids’, ‘Mitochondrial Electron Transport Chain’, ‘Glutathione Metabolism’, and ‘Citric Acid Cycle’, among others (Fig. 5I). In line with these metabolic changes, our proteomic analysis on NDI1-treated PS19 brains highlighted changes to cellular response pathways consistent with NDI1-mediated mitochondrial remodeling and changes to oxidative stress (Fig. S8 A – D). Taken together, these data are highly indicative of NDI1 treatment-associated remodeling of mitochondrial metabolism and highlight extensive downstream changes to both global metabolome and proteome in the PS19 brain.

Since Complex I restoration was only one of the exercise-associated changes in PS19, we wanted to understand similarities between NDI1 treatment vs exercise-associated metabolic changes. Of the 24 rescue metabolites, only 18 were detected in the NDI1 cohort. Of those, 15/18 were significantly changed by NDI1 treatment (Fig. 5J). While not all 15 metabolic changes were directionally conserved with exercise, all MAS-associated metabolites increased by exercise in PS19 were also increased by NDI1 (Fig. 5K). All PS19 brains across exercise and NDI1 cohorts were pooled, and their brain-wide MAS module expression was scored. Just as for exercise, NDI1 treatment restored MAS module scores near WT levels (Fig. 5 L and M). Concordant rescue across all 3 endpoints – tau pathology, spatial metabolome, and brain proteome – establishes augmenting neuronal Complex I activity as sufficient to reproduce the multi-modal exercise-associated phenotype – tau pathology, spatial metabolome, and brain proteome – in PS19 mice.

## Discussion

Collectively, our findings position neuronal Complex I as a convergent bioenergetic axis associated with the exercise training response in primary tauopathy. Independent metabolomic and proteomic signatures converged on the malate–aspartate shuttle and Complex I electron flow. A neuron-targeted genetic bypass of Complex I, in the absence of exercise, reproduced the core anti-tau and metabolic features of the response. Concordance of 3 orthogonal endpoints (anti-tau pathology, spatial brain metabolome, and whole-brain proteome) in the same animals distinguishes augmenting neuronal Complex I activity as sufficient to reproduce the anti-tau and metabolic phenotype seen with exercise, rather than acting as a non-specific correlate. These findings in aggregate suggest that neuronal bioenergetics represent an actionable source of therapeutic targets for tauopathy. Sustained synaptic activity requires continual re-oxidation of cytosolic NADH, and the malate–aspartate shuttle delivers these reducing equivalents to Complex I and the downstream electron transport chain (40, 41). Complex I deficiency, in turn, is the most consistent bioenergetic defect described in AD platelets and post-mortem cortex over the past 3 decades (34–38). Aspartate, glutamate, and malate cycle as shuttle intermediates and constitute the precursor pool for excitatory neurotransmission, a key finding since glutamatergic NMDA receptors have been targeted therapeutically in AD (68, 69). Thus, restoring neuronal Complex I activity may engage this clinically validated axis by maintaining metabolite homeostasis.

Comparing exercise to NDI1 also revealed a shared redox signature. Both interventions raised brain levels of glutathione, ascorbic acid, taurine, and related redox-active species. Because NDI1 transfers electrons from mitochondrial NADH to ubiquinone without proton pumping, it can alter mitochondrial electron handling independently of mammalian Complex I (42, 43). This lowers the proton-motive force that drives reverse electron transport and limits the superoxide leak characteristic of tauopathy (70, 71). Tau hyperphosphorylation and aggregation are themselves redox-sensitive (29), so restoring the antioxidant pool likely feeds back on the pathology. The shared rise in antioxidants with the changes in Complex I subunits and assembly factors is consistent with a unified electron-handling phenotype in which restored forward electron flow lowers redox stress while sustaining ATP supply.

The brain proteomic signature measured came from bulk tissue. Thus, observed Complex I and malate–aspartate shuttle changes could reflect coordinated mitochondrial remodeling in neurons, glia, or even neurovasculature. Neuronal mitochondrial activity is regulated non-cell-autonomously through several well-characterized routes. Astrocytic glutamate uptake activates aerobic glycolysis and lactate release, supplying oxidizable substrate that feeds the neuronal TCA cycle and the malate–aspartate shuttle (71, 72). Astrocytes are the principal source of cysteine and glutathione precursors for neurons, and their supply of these substrates maintains the neuronal redox environment that supports Complex I assembly and activity (73). Astrocytes can also transfer functional mitochondria to neurons via CD38-dependent extracellular release, augmenting the neuronal mitochondrial pool under stress (74). Our observation that neuron-targeted Complex I bypass via NDI1 is sufficient to reproduce the core anti-tau and metabolic features of exercise allows for additional possibilities: 1) an additional exercise-induced glial contribution that exercise engages in parallel and NDI1 addresses only indirectly through restored neuronal energy balance; or 2) non-specific effects due to NDI1 itself.

These findings should be interpreted within the framework established by the human literature: exercise prevents or delays cognitive decline in at-risk populations rather than reversing established AD pathology (16–20). Our PS19 cohort initiates intervention at 3 months, when synaptic mitochondrial tau is already present (27) but late-stage neurodegeneration has not occurred, and is therefore most analogous to early-intervention or secondary-prevention scenarios. The 70%+ reduction in hippocampal phospho-tau immunoreactivity at 6 months is most parsimoniously interpreted as exercise blunting the trajectory of tau pathology accumulation, rather than clearance of established neurofibrillary aggregates. Whether neuronal Complex I restoration carries equivalent benefit at advanced disease stages, when established pathology, BBB compromise, and widespread neurodegeneration are present, remains untested and represents an important translational question.

A second consideration concerns the metabolomic readout. The malate–aspartate shuttle metabolites we identify (aspartate, glutamate, malate, oxaloacetate) are not exclusively committed to feeding NADH equivalents into Complex I; the same molecules participate in TCA cycling, glutamate–glutamine cycling, aspartate-derived biosynthesis, and excitatory neurotransmission. MALDI-MSI measures steady-state metabolite pools rather than flux, and our proteomic data do not establish that the observed metabolite changes preferentially route through Complex I. The convergent metabolomic and proteomic signatures we describe are therefore most accurately framed as engagement of the MAS/Complex I axis rather than direct flux through it.

The exercise–brain field has demonstrated that peripheral exercise-linked signals reach the diseased brain. For example, 1) muscle-derived irisin acts at hippocampal synapses (56, 75–77), with BDNF likely deriving primarily from platelets (78) rather than muscle, 2) liver-derived GPLD1 acts through the cerebrovasculature (79, 80), 3) plasma clusterin dampens neuroinflammation (81), and 4) muscle-derived extracellular vesicles support microglial amyloid clearance (82). Further, exercise drives cortical synaptic protein lactylation in the brain (83), while the MoTrPAC consortium has catalogued these signals across tissues and timepoints (7, 52). These programs have been characterized predominantly through cerebrovascular, glial, or circulating-factor relays before reaching neurons. Our work adds the neuron-intrinsic destination (neuronal Complex I and the malate–aspartate shuttle) as a candidate convergence point for diverse exerkine inputs, allowing the field to ask which combinations of peripheral signals engage this axis and through which downstream cellular intermediates. The accompanying multi-organ spatial atlas (Figs. 1 and 2) is a discovery resource concerning exercise effects across organ systems.

The most immediate open question is: which (upstream) peripheral signal or combination of signals drives the exercise-induced neuronal Complex I and malate–aspartate shuttle alterations? Our multi-organ atlas argues against a single molecule. Brain, lung, and liver covary in their exercise response while muscle and cardiac tissue move in the opposite direction. This complex picture is more consistent with a circulating exercise cocktail vs a single exerkine. Regardless, plasma emerges as the most parsimonious potential integrator and the next therapeutic node to interrogate. Identifying plasma factors that engage this neuronal axis would enable exercise mimetics for patients who cannot exercise. A second open question is the therapeutic window. Tau pathology accumulates over years (23), and whether a point of no return exists beyond which exercise cannot reverse the disease state remains unknown. Testing initiation at later disease stages in PS19 could address this. A third open question concerns translation: whether neuronal Complex I restoration here translates to learning and memory benefits remains to be tested directly.

## Materials and Methods

### Animals

All animal procedures were approved by the University of Florida Institutional Animal Care and Use Committee and were conducted in accordance with the NIH. Hemizygous PS19 transgenic mice (B6;C3-Tg(Prnp-MAPT*P301S)PS19Vle/J; The Jackson Laboratory, stock #008169) and non-transgenic littermate or strain-matched C57BL/6J wild-type (WT) controls (The Jackson Laboratory) were obtained directly from JAX and acclimated to the institutional vivarium for at least one week prior to the start of any intervention. Animals were maintained on a standard (non-reversed) 12 h:12 h light:dark cycle, with lights on at 06:00 and lights off at 18:00 local time, with ad libitum access to standard chow and water. All mice used in this study were single-housed beginning at the onset of intervention (3 months of age) to enable individual cage-level activity quantification and to standardize housing between sedentary (control) and wheel-running cohorts. Animal health and cage conditions were monitored daily by trained laboratory personnel in addition to vivarium staff, and home-cage activity was continuously recorded with cage-top infrared activity monitors (Actimetrics ClockLab Wireless Node) placed on top of every cage for the full duration of the intervention. For the voluntary wheel-running arm specifically, in-cage wheel revolutions were additionally recorded continuously via ClockLab (Actimetrics) as described below (see “Voluntary wheel running”). Both the voluntary wheel-running cohort (WT and PS19, sedentary and exercise) and the AAV9 ICV cohorts (AAV9-Control and AAV9-SYN1-NDI1) were composed exclusively of female mice to minimize sex-as-a-biological-variable noise across the samples. Animals were assigned to experimental groups using a randomization scheme controlling for body weight, and experimenters performing surgery, tissue collection, imaging acquisition, and image analysis were blinded to group identity wherever feasible.

### Experimental design and group structure

This study was carried out in two sequential, conceptually linked arms, both performed on a tauopathy-relevant timeline (3 to 6 months of age). The voluntary wheel-running (VWR) cohort (Arm 1) was performed first, and the metabolomic and proteomic observations from this cohort directly motivated the subsequent intracerebroventricular (ICV) AAV9-NDI1 intervention (Arm 2). Arm 2 was therefore designed as a targeted genetic test of whether augmenting neuronal NADH dehydrogenase activity is sufficient to recapitulate the anti-tau effects observed in Arm 1.

### Arm 1: Voluntary wheel running

The VWR experiment was a WT vs PS19 × sedentary vs exercise full factorial groups: sedentary WT, exercise WT, sedentary PS19, and exercise PS19 (n = 3 mice per group). All animals were enrolled at 3 months of age, with WT and PS19 mice purchased directly from The Jackson Laboratory and acclimated to the vivarium before the start of the intervention. Exercise animals were single-housed in voluntary wheel-running cages as described previously (84) with continual, ad libitum access to an in-cage running wheel from 3 to 6 months of age. Sedentary controls (sedentary WT and sedentary PS19) were single-housed in matched cages without a wheel for the same duration. Three days prior to takedown, running wheels were mechanically locked in place to eliminate acute exercise effects on the brain metabolome and proteome. Cage-top activity monitors recorded ambulatory activity continuously across all four groups, and animal welfare was inspected daily by laboratory personnel.

### Arm 2: Bypass of mitochondrial Complex I (AAV9-NDI1)

PS19 mice (n = 3 per group) received a single ICV injection at 3 months of age of either: an empty vector control (ultra-purified AAV9-EGFP, parental vector VB010000-9287ffw; “AAV9-Control”) or (ii) a neuron-restricted NDI1 vector under the human synapsin I promoter (pAAV[Exp]SYN1>sce_NDI1[NM_001182483.1](ns):P2A:EGFP:WPRE; VectorBuilder VB250224-1374sqf; “AAV9-SYN1-NDI1”). After surgery, animals were returned to their home cages and aged undisturbed until 6 months of age, at which point they were taken down for tissue collection.

### Voluntary wheel running

Beginning at 3 months of age, WT and PS19 mice randomized to the exercise condition were single-housed in voluntary wheel-running cages consisting of a stainless-steel running wheel (11 cm inside diameter, 5.4 cm wide; 1.2 mm wide bars spaced 7.5 mm apart). The wheel was supported by a shaft running through its center that rests in holes drilled in each side of the cage bottom; a modified wing nut mounted on the end of the shaft served as the counting trigger, striking an external electromechanical switch on every revolution. This wheel-cage configuration is currently distributed by Actimetrics as the model PT2-MCR1 wheel-running cage. Sedentary controls were single-housed in matched cages without a wheel for the same duration. Mice had continuous, ad libitum access to the wheel for 3 months (3-6 months of age). Each wheel switch was interfaced via the supplied data cable to a dedicated computer, and revolution counts were recorded continuously in 1-min bins using ClockLab acquisition software. Three days before sacrifice, running wheels were physically locked in place to prevent rotation while remaining in the cage, ensuring that any observed brain phenotypes reflect chronic training adaptations vs acute exercise effects. Mice were inspected daily for general health and grooming status by trained laboratory personnel, and bedding was changed biweekly.

### Viral vectors

AAV9 vectors were custom-cloned and packaged by VectorBuilder Inc. The neuronal vector carried a 469 bp human synapsin I (SYN1) promoter driving the Saccharomyces cerevisiae NDI1 coding sequence (NM_001182483.1, 1,539 bp) followed by a porcine teschovirus-1 P2A self-cleaving linker (66 bp) and an EGFP reporter (720 bp), with a WPRE post-transcriptional regulatory element (598 bp) and a BGH polyadenylation signal; viral genome size, 3,984 bp; full vector size, 6,603 bp. The viral control was the ultra-purified EGFP AAV9 supplied by VectorBuilder from parental vector at matched titer. All viral aliquots were stored at -80 °C, thawed on ice immediately prior to use, and kept on ice during the surgical session; aliquots were used undiluted (neat) for ICV delivery.

### Intracerebroventricular (ICV) AAV delivery

At 3 months of age, mice were anesthetized with isoflurane induction, placed in a stereotaxic frame, and prepared for aseptic surgery. Pre-operative analgesia and ophthalmic ointment were administered, and body temperature was maintained on a feedback-controlled heating pad. The scalp was shaved, prepped sequentially with povidone-iodine and 70% ethanol, and a midline incision was made to expose bregma. A small craniotomy was drilled over each lateral ventricle. Bilateral ICV injections were targeted to the lateral ventricles using the following coordinates relative to bregma: AP −0.56 mm, ML ±1.10 mm, DV −2.26 mm from the skull surface. A total of 2 µL of undiluted AAV9 viral stock was delivered per ventricle through a 10 µL Hamilton syringe using an automated microinjector. The needle was held in place for 10 min following completion of each injection to permit diffusion and minimize backflow before slow withdrawal over 5 min. Post-operative monitoring and analgesia were continued for at least 72 h. Mice were aged in their home cages until 6 months of age, when their tissues were harvested.

### Tissue collection and cryosectioning

At 6 months of age, mice were euthanized by cervical dislocation without prior perfusion to preserve metabolite integrity for MALDI-MSI. Brains were rapidly dissected, briefly rinsed in PBS, followed by two water washes, blotted, and flash-frozen over liquid-nitrogen before storage at -80°C. For both MALDI-MSI and immunofluorescence (IF), 10 µm sagittal sections were collected on a Leica CM1860 cryostat at -20°C and thaw-mounted onto standard microscope charged slides. Slides were vacuum-sealed and stored at -80°C until use.

### Immunofluorescence Staining

Fresh-frozen 10 µm sagittal sections were brought from -80°C to room temperature and fixed in freshly prepared 1% paraformaldehyde (PFA; diluted in 1× PBS from 16% PFA, Electron Microscopy Sciences #15710) for 30 min at room temperature. After fixation, slides were washed once in 1× PBS for 10 min, then rehydrated through a graded ethanol series in glass Coplin staining jars (two × 2 min in 100% ethanol; 3 min each in 95%, 70%, and 50% ethanol) and washed once in 1× Tris-buffered saline (TBS) for 10 min. Heat-induced antigen retrieval was performed by immersing slides in 1× citrate buffer (diluted from 10× stock) in a sealed side-opening mailer placed in a water bath at 96°C for 1 h, followed by 30 min cooling at room temperature; slides were then washed once in 1× TBS for 10 min. Tissue sections were outlined with a hydrophobic PAP barrier pen, and blocked for 1 h at room temperature with 200 µL/section of tissue blocking buffer (TBB; 2% v/v normal goat serum and 5% w/v bovine serum albumin in TBS containing 0.05% w/v octyl β-D-glucopyranoside [TBS-OG; Sigma #03757]).

Primary antibodies were diluted in TBB, applied to sections at 200 µL/section, and incubated overnight at 4°C in a humidified, light-protected staining tray. Primary antibodies used in this study were: mouse monoclonal anti-phospho-Tau (Thr231), clone AT180 (Thermo Fisher/Invitrogen #MN1040; dilution 1:300) (85), rabbit polyclonal anti-NDUFV2 (Proteintech, #15301-1-AP; dilution 1:100) (86), rabbit anti-NeuN (for viral cohort, Fig. 5) (abcam, #ab177487) (87), and mouse anti-NeuN (for Fig. S6) (abcam, #ab279295) (88). The following day, slides were washed three times in 1× TBS (5 min each, 800 µL/section) and incubated with species-matched fluorophore-conjugated secondary antibodies, as appropriate for each experiment (donkey anti-mouse Alexa Fluor 488, anti-rabbit Alexa Fluor 488, anti-rabbit Alexa Fluor 594, and anti-mouse Alexa Fluor 647, respectively) diluted in TBB for 1 h at room temperature in the dark. Slides were then washed three times in 1× TBS (5 min each), counterstained with DAPI diluted 1:1000 in 1× TBS for 10 min at room temperature, washed three more times in 1× TBS (5 min each), and coverslipped with ProLong Diamond without DAPI. For the viral cohorts, AAV9-EGFP expression was visualized directly from native EGFP fluorescence in the green channel (488), and the phospho-Tau secondary antibody was replaced with a 647 channel (donkey anti-mouse Alexa Fluor 647).

Whole-section images were acquired on an Olympus VS200 using matched acquisition settings across all samples within a staining batch. Quantitative analysis of hippocampal AT180 immunoreactivity was performed in HALO (Indica Labs) software on operator-drawn hippocampal regions of interest by an experimenter blinded to genotype and treatment. Per-animal hippocampal AT180 measurements were compared between groups performed in GraphPad Prism by unpaired two-tailed Welch’s t-test for two-group, with significance set at α = 0.05. All bar/dot plots show individual animal values as dots, with the mean ± SEM.

### MALDI mass spectrometry imaging (MALDI-MSI)

Fresh-frozen brains were cryosectioned at 10 µm on a Leica CM1860 cryostat at -20 °C. Sagittal sections were thaw-mounted onto standard microscope charged slides, vacuum-sealed, and stored at -80 °C until analysis. Immediately prior to matrix application, slides were brought to room temperature and dried by vacuum desiccation for 1 h to ensure surface dehydration and preserve metabolite/lipid integrity.

### Matrix application

N-(1-naphthyl)ethylenediamine dihydrochloride (NEDC) matrix was applied with an HTX M5 TM-Sprayer (HTX Imaging) using a 7 mg/mL NEDC solution in 70% methanol delivered at 14 passes, 30°C nozzle temperature, 0.06 mL/min flow rate, 10 psi nitrogen pressure, and a 50°C tray temperature.

### Acquisition on the MALDI-timsTOF

Data were collected on a timsTOF fleX mass spectrometer (Bruker Daltonics) equipped with a 10 kHz SmartBeam 3D laser. Region definition and instrument setup were automated using the autopilot feature in flexImaging v6.0; high-resolution optical scans (.tif) were used to generate a standardized geometry (flexImaging .mis) for consistent region masking and pixel alignment across runs. Imaging was performed in negative ion mode with a 46µm laser raster yielding a 50µm pixel pitch. For metabolomics, MS1 spectra were collected from m/z 20-750 at 80% laser power (1 burst of 396 shots) with global instrument settings of 30V MALDI plate offset, -60V deflection 1 delta, and 200Vpp for Funnels 1 and 2 and the multipole RF; collision energy was 7eV with a 700Vpp collision RF.

### MALDI-MSI data processing, quality control (QC), and normalization

Data were imported into SCiLS Lab (2026a Pro, 14.00.17781) for peak picking, alignment, and annotation. Peak annotation was performed against established in-house spectral library. Per pixel data feature intensities were then imported into R (v4.4.3). Ion intensities were normalized at the per pixel level by total ion count (TIC). Next, to reduce technical artifacts (such as outlier pixels), we performed median absolute deviation (MAD) filtering on the TIC-normalized per pixel data (medians were calculated per tissue section). Pixels exceeding +/- 4 MAD from section median were designated as outliers and removed on a per-feature basis. Per pixel data were then mean aggregated (per animal) by either (11) organ, (12) organ-specific metabolic zone (see ‘unsupervised integrated spatial clustering of metabolic zones’), or (1) brain region (see ‘semi-unsupervised brain region identification’). Mean aggregate intensities were multiplied by a constant scaling factor and subsequently log2-transformed. Lastly, to correct for batch-to-batch technical variability, we performed batch correction across organs using the ComBat empirical Bayes framework (‘sva’ package, v3.54.0). Replicate was modeled as the batch variable. Treatment (and genotype, as appropriate) were included as biological covariates in the ComBat design matrix to preserve treatment-, and genotype-, associated variation during batch correction. These data were used for downstream differential expression analysis.

### Unsupervised integrated spatial clustering of metabolic zones

To identify unified metabolic zones (clusters) per organ, molecular features from individual tissue sections were integrated using linear batch harmonization implemented in removeBatchEffect (limma, v3.60.6), treating section as the integration variable (performed separately per organ). Principal component analysis (PCA) (prcomp function, base R) was then performed, and the top 20 PCs were retained. Optimal cluster number was then determined for each organ using the Calinski-Harabasz (C-H) criterion by evaluating k-means solutions (k = 2-10) on a random subset of 5,000 pixels. The cluster number maximizing the C-H index was selected, and final k-means clustering was performed on the full PCA embedding. For visualization, UMAP (uwot package, v0.2.4) was applied to the integrated feature matrix.

### Semi-unsupervised brain region identification

Brain regions were identified using a graph-based clustering workflow, followed by manual pixel annotation clean up. To do this, a representative wild-type brain section was selected as a reference. Features were filtered to retain metabolites detected >80% of pixels. Then, features were ranked by MAD, and the 600 most variable features were selected. PCA (prcomp) was performed using these variable features, and the top 30 PCs were retained. Two independent k-nearest-neighbor graphs were then constructed (RANN, v2.6.2), representing spatial proximity and feature similarity. These graphs were integrated into a single spatial-molecular network, and community detection was performed using the Leiden algorithm (leidenbase, v0.1.36), with Louvain clustering (igraph, v2.3.1), where necessary. Multiple clustering resolutions were evaluated to identify most anatomically coherent region domains, resolution = 2.4 was used. To assign integrated region annotations across brains, all sections were integrated in PCA space using Harmony (harmony, v2.0.2), with animal identity used as batch variable. Representative section-derived region centroids were then used to assign pixels from each animal to the nearest reference region in the integrated space using nearest-neighbor classification (FNN, v1.1.4.1). Region labels were spatially smoothed within each animal to reduce isolated pixel-level artifacts. Lastly, the resulting region maps were manually reviewed and corrected, as needed, to ensure appropriate region borders.

### MALDI-MSI differential expression (DE) analysis

Differential expression (DE) analyses were performed on mean aggregate data using the limma framework. Feature-wise linear models were fit using empirical Bayes variance moderation, and DE statistics were calculated from predefined contrasts using moderated t-tests. Model designs were specified according to the biological comparisons, including genotype, treatment, anatomical region/cluster, and interaction terms where appropriate. Whole organ-specific DE analysis (exercise): DE analyses were performed independently within each organ. For each organ, feature abundances were modeled as a function of treatment using a linear model (∼ treatment), and the exercise vs control contrast was evaluated using empirical Bayes-moderated statistics. Brain region-specific DE analysis: DE analyses were performed independently for each annotated brain region. For each region, molecular abundances were modeled using a factorial design including genotype, treatment, and genotype-by-treatment interaction terms (∼ genotype x treatment). Contrasts were used to estimate exercise effects within genotypes, genotype effects within treatment group, and genotype-by-treatment interaction effects. Integrate brain-wide DE analyses: To identify conserved changes across brain regions, while accounting for regional structure, all annotated brain regions were modeled jointly using a design matrix including experimental group (e.g., control vs NDI1 or WT+CTRL vs WT+Ex vs PS19+CTRL vs PS19+Ex) and region as fixed effects (∼ 0 + group + region). Since multiple region-level measurements were obtained from each animal, within-animal correlation was estimated using duplicateCorrelation (limma), and animal identity was included as a repeated-measures blocking factor. Contrasts were used to compare genotype under control conditions and to estimate treatment-associated changes within PS19 genotype.

### Inter-organ correlation analysis

To compare the exercise-associated metabolic responses across organs, Spearman correlation was calculated across organ pairs using log2(fold change) estimates from organ-specific DE analysis. Only features detected across all organs were used. Pairwise similarity was visualized as a clustered correlation heatmap using average-linked hierarchical clustering.

### Whole organ metabolic pathway analysis

Metabolic pathway analysis was performed using MetaboAnalyst (v6.0) (89). For each organ-specific DE analysis, differentially expressed features (FDR < 0.1) were used as input (query). Significant features were mapped to pathways using the mus musculus Small Molecular Pathway Database (SMPDB). Background features were restricted to features detected within that organ. Analysis was performed using default parameters.

### Brain region metabolites set enrichment analysis (MSEA)

Brain region-level MSEA was performed using fgsea (v1.30.0). For each region, features were ranked by limma-derived log2(fold change) and pre-ranked analysis was performed against SMPDB pathway feature sets (restricted to detected features). Pathways were restricted to curated SMPDB terms represented by >3 features. Normalized enrichment scores (NES) were used as pathway-level scores, and pathways were summarized across regions by the number of regions reaching FDR < 0.1 and mean absolute NES.

### Over-representation analysis (ORA) of rescue metabolites

ORA was performed using ‘Enrichment Analysis’ within MetaboAnalyst (v6.0). For exercise rescue in PS19, the query features were selected by meeting the following criteria: increased by exercise in PS19 (PS19+Ex vs PS19+Ctrl, FDR < 0.1, log2FC > 0) and decreased by PS19 (PS19+Ctrl vs WT+Ctrl, log2FC < 0). Significant features were mapped to pathways using the *mus musculus* Small Molecular Pathway Database (SMPDB). Background features were restricted to features detected within that organ. Analysis was performed using default parameters.

### Malate-aspartate shuttle (MAS) module scoring and visualization

Malate-aspartate shuttle (MAS) module was comprised of the following features: malic acid, aspartic acid, oxaloacetic acid, and glutamic acid. To calculate module score, all detected features were grouped into 24 bins based on their average intensity across pixels. For each module feature, 25 abundance-matched control features were randomly selected from the same bin. The module score for each pixel was then calculated as the average abundance of the module features minus the average abundance of the matched control features, controlling for differences in feature abundance distributions across the dataset. For representative MAS modules scores in Fig. 3, animal-level module scores were normalized relative to the cohort-specific PS19 control group (either sedentary PS19 animals or control AAV-treated PS19 animals) by subtracting the control-group mean and dividing by the control-group standard deviation. Consequently, a normalized score of 0 = mean PS19 control animal for that cohort, whereas values of ±1 represent one control-animal SD above or below the cohort-specific control mean.

### Global proteomic sample preparation

For whole-brain proteomics, mice (n=3 per group) were euthanized by cervical dislocation, brains were rapidly dissected, briefly rinsed in PBS then water, and flash frozen over liquid-nitrogen. Hemi-brain samples were then used for downstream protein extraction. Brain proteins were extracted and digested using EasyPep™ MS Sample Prep Kit (Thermo Fisher Scientific). The total protein concentration for each sample was determined on a Qubit fluorometer (Invitrogen), and appropriate volume was collected to equal 20 µg total protein for digestion. Following digestion using sequencing-grade trypsin/Lys-C (Promega Rapid Digestion Kit) according to the manufacturer’s recommendations. Three sample volumes of rapid digestion buffer were added, samples were incubated with 1 µL of 0.1 M dithiothreitol in 100 mM ammonium bicarbonate at 56 °C for 30 min, alkylated with 0.54 µL of 55 mM iodoacetamide in 100 mM ammonium bicarbonate at room temperature in the dark for 30 min, and digested with 1 µL of freshly prepared trypsin/Lys-C (1 µg/µL in rapid digestion buffer) at 70 °C for 1 h. The digestion was quenched with 0.5% trifluoroacetic acid (TFA), and tryptic peptides were analyzed by LC-MS/MS immediately to ensure high-quality tryptic peptides with minimal non-specific cleavage.

### Nano-LC-MS/MS Acquisition

Nano-liquid chromatography tandem mass spectrometry (nano-LC/MS/MS) was performed on a Thermo Scientific Q Exactive HF Orbitrap mass spectrometer equipped with an EASY-Spray nanospray source (Thermo Scientific) operated in positive ion mode. The LC system was an UltiMate™ 3000 RSLCnano (Thermo Scientific). Mobile phase A was water with 0.1% formic acid and mobile phase B was acetonitrile with 0.1% formic acid; the loading-pump mobile phase A was water with 0.1% trifluoroacetic acid. Then, 5 µL of each sample were injected onto a Thermo Scientific µPAC™ C18 trapping column (C18, 5 µm pillar diameter, 10 mm length, 2.5 µm inter-pillar distance) at 10 µL/min, held for 3 min, and washed with 1% B to desalt and concentrate the peptides before switching to inject. Peptides were eluted from the trap onto a Thermo Scientific 110-cm mPAC analytical column (C18, 5 µm pillar diameter, 110 cm length, 2.5 µm inter-pillar distance) maintained at 40°C, with a flow rate of 750 nL/min for the first 15 min and then reduced to 250 nL/min. Peptides were separated into the Q Exactve System using a gradient of 1% B to 20% B over 100 min and then to 45% B over 20 min, for a total LC run time of 150 min. The EASY-Spray source operated at 1.5 kV with a capillary temperature of 200°C. MS/MS was acquired using the TopTenTM method: a full MS1 scan from m/z 375-1575 Da at 60,000 resolution and MS2 scans at 15,000 resolution on the fifteen most abundant precursors to define the amino acid sequence. AGC targets were 3x106 (MS1) and 2x105 (MS2); maximum injection times were 50 ms (MS1) and 55 ms (MS2); microscans were set to 1 for both. HCD fragmentation used a stepped (N)CE of 28 with an isolation window of 4 m/z. Singly charged ions were excluded from MS2 selection, dynamic exclusion was enabled with a repeat count of 1 within 15s and isotopes excluded, and a siloxane background peak at m/z 445.12003 was used as the internal lock mass. Instrument performance was monitored between batches using a HeLa protein digest standard with columns replaced and the instrument cleaned whenever HeLa protein IDs fell below 3,200.

### Database Search and Label-Free Quantitation

All MS/MS spectra were searched using the Chimerys node (MSAID; Thermo Fisher Scientific, San Jose, CA, USA) in Proteome Discoverer v3.2.0.450. The search database was the Mus musculus reference proteome from UniProt (sp_incl_isoforms, TaxID = 10088 and sub-taxonomies, Release 406, v2024-10-02) concatenated with the 0602 Universal Protein Contaminants library (Frankenfield et al., J. Proteome Res. 21, 2104–2113, 2022; “0602_Universal Contaminants.fasta”) with the assumption of digestion enzyme trypsin. Chimerys was run with the inferys_4.7.0_fragmentation prediction model, a precursor mass tolerance of 10.0 ppm, and a fragment ion mass tolerance of 0.020 Da. The single static modification was carbamidomethylation of cysteine (Carbamidomethyl, C). Dynamic modifications were oxidation of methionine (Oxidation, M) and phosphorylation of serine, threonine, and tyrosine (Phospho, S; Phospho, T; Phospho, Y), with a maximum of three dynamic modifications per peptide. Precursor-ion-intensity label-free quantitation (LFQ) was performed in Proteome Discoverer (Thermo Fisher Scientific v3.2.0.450). Groups for Arm 1 and 2 were compared using a non-nested study factor (Arm 1: genotype × condition; Arm 2: viral group), normalization was determined using all peptides, and protein abundances were calculated by summed abundances of the connected peptide groups.

### Proteomic DE analysis

Proteomic DE analysis was performed using the limma framework. Protein abundances were log2-transformed and proteins detected in fewer than eight samples were excluded from downstream analysis. Samples were modeled as four experimental groups (WT+Ctrl, WT+Ex, PS19+Ctrl, and PS19+Ex), and empirical Bayes variance moderation was performed with trend and robust estimation. Predefined contrasts were used to estimate the exercise effect in WT animals (WT+Ex vs WT+Ctrl), the exercise effect in PS19 animals (PS19+Ex vs PS19+Ctrl), the genotype effect under baseline conditions (PS19+Ctrl vs WT+Ctrl), and the genotype effect following exercise (PS19+Ex vs WT+Ex). Statistical significance was assessed using moderated t-tests, and multiple-testing correction was performed using the Benjamini-Hochberg procedure. For NDI1 treatment proteomics, DE analysis was performed similarly, using only two experimental groups (PS19+Ctrl AAV and PS19+NDI1 AAV).

### Weighted protein co-expression network analysis

Weighted gene co-expression network analysis (WGCNA) was performed on the bulk brain proteomics dataset using the WGCNA package (v1.72.5). Proteins detected in fewer than eight samples were excluded, and sample and protein quality were assessed using ‘goodSamplesGenes’. A signed protein co-expression network was constructed using bi-weight midcorrelation with pairwise complete observations and a maximum outlier proportion of 0.1. The soft-thresholding power was selected as the first power achieving a signed scale-free topology fit index of R² ≥ 0.80; if no power met this criterion, a power of 8 was used. Protein modules were identified using ‘blockwiseModules’ with a signed topological overlap matrix, deepSplit = 2, minimum module size of 30 proteins, and module eigengene merge cut height of 0.25. Module eigengenes were calculated as the first principal component of each module and correlated with experimental traits, including genotype, treatment, genotype-by-treatment interaction, and four-level experimental group identity (WT+Ctrl, WT+Ex, PS19+Ctrl, and PS19+Ex). Module-trait associations were assessed using Pearson correlation, and associated P values were calculated using Student asymptotic tests. Module membership was calculated as the correlation between each protein and module eigengenes, and gene significance was calculated as the correlation between each protein and the exercise trait.

### Module eigengene relationship analysis

Relationships among WGCNA modules were assessed by calculating pairwise Pearson correlations between module eigengenes. Statistical significance of module-module correlations was evaluated using two-sided Pearson correlation tests and resulting P values were corrected for multiple testing using the Benjamini-Hochberg false discovery rate procedure. To visualize module organization with respect to experimental structure, mean module eigengene values were calculated for each experimental group, and group means were z-scored within each module to highlight relative module enrichment across WT+Ctrl, WT+Ex, PS19+Ctrl, and PS19+Ex groups. Module eigengene relationships were visualized using pheatmap (v1.0.12).

### Anti-module axis analysis

Because the brown and red modules exhibited a significant inverse eigengene correlation, an anti-module axis was defined as the standardized difference between their eigengenes, z(ME_brown − ME_red), to capture the dominant pattern of reciprocal module activity across samples. For each sample, the axis was calculated as the standardized difference between the brown and red module eigengenes, z(ME_brown − ME_red), such that positive values reflected a brown-dominant module state and negative values reflected a red-dominant module state. Protein-level axis scores were calculated as the Pearson correlation between protein abundance and the anti-module axis across samples. Module hubness was quantified as the maximum absolute module membership for the brown module, max(|kME_MEbrown|) Candidate proteins associated with the anti-module axis were identified using thresholds of |axis score| ≥ 0.8 and hubness ≥ 0.8.

### Proteomic pathway analysis

Pathway analysis of significantly altered proteins was performed within EnrichR (90), against the Gene ontology (GO) Biological Process 2026 database. Background proteins were restricted to all detected proteins in our proteomic dataset. Query proteins for exercise-associated rescue pathway analysis were defined as proteins significantly altered in PS19 control mice relative to WT controls (FDR < 0.05) and significantly altered by exercise within PS19 mice (FDR < 0.05), where exercise-induced changes occurred in the opposite direction of genotype-associated changes (opposite-signed log2 fold-changes), consistent with a rescue-like response. Query proteins for NDI1 treatment-associated pathway analysis were defined as proteins significantly altered by NDI1 treatment relative to control AAV treatment (FDR < 0.05).

### Complex I component mRNA expression via MERFISH

Publicly available MERFISH data from the adult mouse brain atlas generated by Zhang et al. (Nature, 2023) (67) were accessed through the CELLxGENE Explorer platform. Spatial expression maps and UMAP embeddings were generated for individual genes encoding mitochondrial electron transport chain complex I subunits. PNG images of UMAP embeddings and corresponding spatial expression maps were exported directly from the browser interface. Gene sets consisted of exercise-associated complex I proteins identified in the PS19 exercise proteomic analysis. The CELLxGENE gene set tool was used to generate composite expression score spatial expression maps for these genes. Resulting visualizations were used for qualitative assessment of cell type-specific and region-specific enrichment patterns across the adult mouse brain.

## Acknowledgements

This study was supported by National Institutes of Health (NIH) grants R01AG066653, R01CA266004, R01AG078702, R01CA288696, and RM1NS133593 to R.C.S., R35NS116824 to M.S.G., and R01HL149800 to G.S.M. Z.L. is supported by the MBI Gator NeuroScholar Program. S.Q. is supported by the MBI Gator NeuroScholar Program in partnership with the Nancy A. Fackler Endowment for Brain Research, as well as NIH T32 CA257923 and P30 CA247796. T.M. is supported by NIH T32 HL134621. C.S. is supported by the NMPT program under NIH T32 HD043730. Additional support was provided by the McKnight Brain Institute and the Center for Advanced Spatial Biomolecule Research (CASBR) at the University of Florida College of Medicine. We thank Manuel Sanchez for technical support. All animal procedures were approved by the University of Florida Institutional Animal Care and Use Committee (IACUC). Large language models (Claude) were used for grammar checking and proofreading.

## Author Contributions

Conceptualization, R.C.S. and K.A.E.; Methodology, R.C.S., X.M., M.S.G., and C.W.V.K.; Formal Analysis, T.M., S.Q., Z.L., X.M., and S.F.; Investigation, T.M., S.Q., Z.L., L.W., B.Z., C.S., X.M., A.R., R.L., C.M.S., V.B-C., A.T., K.W., N.R., and S.F.; Resources, M.A.G-M., G.S.M., C.W.V.K., M.S.G., N.S.C., and K.A.E.; Writing – Original Draft, R.C.S.; Writing – Review & Editing, R.C.S., N.S.C., M.S.G., C.W.V.K., G.S.M., and K.A.E.; Funding Acquisition, R.C.S., M.S.G., N.S.C., and K.A.E.; Supervision, R.C.S., M.S.G., and K.A.E.

## Data and Materials Availability

Spatial metabolomics data generated in this study will be available from sunlabresources.com upon acceptance. Proteomics data will be deposited in the ProteomeXchange database. No new code was produced for this study.

